# Saving water to get ‘more crop per drop’ – A new phenotyping framework revealed wide plasticity in wheat

**DOI:** 10.1101/2025.08.13.670015

**Authors:** Brian Collins, Karine Chenu

## Abstract

Improving transpiration efficiency (TE) offers a pathway to increase yield in drought-prone environments. This study examined genotypic variation in TE and its physiological determinants across diverse wheat lines. An initial experiment with six cultivars was expanded to 105 genetically-diverse genotypes evaluated under well-watered conditions and fluctuating vapour pressure deficit (VPD). Using a high-throughput lysimeter platform, transpiration rates were recorded every 10 minutes and normalised daily at low VPD to account for genotypic variations in canopy size. TE was strongly associated with reduced normalised transpiration rate at high VPD (TR_norm-highVPD_), while no significant relationship was found with maximum photosynthetic capacity. High-TE lines achieved either greater biomass with similar water use or similar biomass with lower water use, reflecting a water-saving strategy under high evaporative demand. A complementary experiment under low VPD revealed limited genotypic variation in intrinsic TE, reinforcing the value of TR_norm-highVPD_ as a screening trait. Consistent correlations between TE and TR_norm-highVPD_ across experiments highlight the potential of TR_norm-highVPD_ as a robust phenotyping target. Several high-TE lines outperformed modern cultivars, offering promising sources of novel alleles. These findings provide a scalable framework to identify drought-resilient genotypes and support breeding strategies to improve water productivity and achieve more crop per drop.

**HIGHLIGHT:** Wheat genotypes with reduced normalised transpiration rate under high evaporative demand achieved greater transpiration efficiency. A new automatised high-throughput phenotyping framework is proposed to assist drought-resilient breeding.

## 1 INTRODUCTION

A common challenge faced by wheat growers and breeders in rain-fed cropping systems is the lack of plant available moisture, which often limits crop production (Araus et al., 2002). Increased transpiration efficiency (TE) can potentially assist breeders selecting for germplasm that produces ‘more crop per drop’ to increase grain yield in water-limited environments (Condon et al., 2002; Marris, 2008; Chenu and Fletcher, 2015; Chenu et al., 2018; Christy et al., 2018; Vadez et al., 2024).

In the Australian wheatbelt, wheat production is severely constrained by drought and the pronounced variability of annual rainfall (Murphy and Timbal, 2008; Chenu et al., 2011, 2013; Rebetzke et al., 2013). Over the past three decades, this challenge has been further intensified by a marked increase in drought and heat stress, greater frequency of concurrent temperature– precipitation extremes (Ababaei and Chenu, 2020; Collins and Chenu, 2022; Collins, 2022) and exponential rise in the atmospheric vapour pressure deficit (VPD; Hersbach et al., 2020). In these environments, TE has been found as a trait highly impacting wheat productivity in simulation studies (Casadebaig et al., 2016), especially where crops rely heavily on stored soil water (Condon et al., 2002; Ababaei and Chenu, 2019a; Collins et al., 2021). Experimental work, based on a surrogate trait for TE, has also highlighted the potential of TE to increase yield in crops like wheat (Rebetzke et al., 2002, 2009). Interestingly, TE has significantly increased with breeding in Australian cultivars over recent decades (Chenu and Fletcher, 2015; Fletcher and Chenu, 2015). Further improvements may be raised by better understanding the underlying processes and their quantitative contribution to TE, and by importing new alleles to breeding germplasm pools.

Typically defined at the plant level, TE corresponds to the amount of biomass produced per unit of water transpired. TE is generally measured in sealed containers excluding soil evaporation and drainage and differs from ‘water use efficiency’ (WUE) that commonly refers to field measurements, which include soil evaporation and drainage (Richards et al., 2002). TE at the plant level is typically measured over a period of days to weeks and provides limited insights into the quantitative response to environmental conditions. However, many of physiological processes determining TE operate at the leaf level, at which TE can be defined as the ratio between assimilation (photosynthesis) and the flux of water vapour through the stomata, measured over a period of seconds to minutes.

TE components of interest include (i) photosynthetic capacity, (ii) the ability to limit transpiration rates per unit leaf area under high VPD, and (iii) the threshold soil water content at which transpiration rates respond to soil drying (Chenu et al., 2018). In crops like wheat, genetic variations have been identified for photosynthetic capacity and efficiency (e.g. Silva-Pérez et al., 2020), which can now be measured in the field at high-throughput levels with hyperspectral reflectance (Silva-Pérez et al., 2020). Wheat genotypes also vary for their ability to limit transpiration rates per unit leaf area under high evaporative demand (Schoppach and Sadok, 2012; Schoppach et al., 2016, 2017), and under low soil water content (Schoppach and Sadok, 2012). Phenotyping of these later traits is typically done in controlled environments and has traditionally required substantial labour resources with frequent measurements of both water use and leaf area development.

While genotypic variations have been identified for TE component traits, their direct contribution to genotypic variation in the plant-level TE has not, to our knowledge, been quantified experimentally. Simulations studies have however established the importance of limiting maximum transpiration rates to increase TE and ultimately grain yield under drought (e.g. Collins et al., 2021). While transpiration rate generally increases with increasing VPD, TE declines with increasing VPD (Kemanian et al., 2005). Limiting maximum transpiration rates, i.e. reducing transpiration at high VPD, corresponds to a reduction in transpiration when water use is least efficient and TE is lowest. This water-saving trait reduces the water use and increases the efficiency of biomass produced per unit of water (TE). In addition, a reduction in pre-anthesis water use due to such limited maximum transpiration rate can delay or reduce a water stress later in the crop cycle, especially where crops rely on stored soil moisture (Collins et al., 2021; Borrell et al., 2023). *In silico* studies for sorghum, soybean and maize in different cropping area have shown that limiting maximum transpiration rates generally increases grain productivity in regions prone to post-flowering drought (Sinclair et al., 2005, 2010; Messina et al., 2015). Under well-watered conditions however, restricting water use at high VPD due to reduced stomatal conductance can restrict CO_2_ assimilation and result in yield loss (Sinclair et al., 2005; Messina et al., 2015; Ababaei and Chenu, 2019a; Collins et al., 2021). Collins et al. (2021) proposed a model in wheat to limit transpiration rate proportionally to VPD above a threshold, as observed experimentally (rather than capping maximum transpiration rate to a fixed level as previously done). They found that reducing transpiration rates at high VPD resulted in greater TE and grain yield across drought-prone Australian environments, especially where crops heavily rely on stored soil water, and with increasing benefits in future climate scenarios.

This study aimed to understand processes underlying genotypic variations in TE in wheat and to identify promising lines with high TE. To do so, a few wheat genotypes contrasting in TE were grown under well-watered conditions (i) in a polyhouse with variable VPD over two seasons (2018-2019) and (ii) under low VPD in a growth room to measure intrinsic TE. In addition, a wide range of genetically diverse wheat accessions was phenotyped along with elite lines to explore options to increase TE and its components in the Australian wheat germplasm pool.

## 2 MATERIALS AND METHODS

### 2.1 Experiments with fluctuating evaporative demand

Two experiments were conducted in a high-throughput automated lysimeter platform (Fig. 1, Table 1) in 2018 (Exp 1) and 2019 (Exp 2). The system, as described by Chenu et al. (2018), combines the concept of a constant water table (Hunter et al., 2012) with automatic monitoring and irrigation. The platform, located at Gatton, Queensland (27°32’19 S, 152°20’04 E) in a solar weave tunnel, consists of 560 lysimeters (4L ANOVApot®, Anova Solutions). Watering and weighing were performed fully automatically every 10 min and average weight of each lysimeter since the last reading was recorded. A weather station located in the centre of the tunnel monitored environmental conditions at 10-min intervals. Four plants per pots were sown 3 cm deep in a potting media with 2.8 g L^-1^ of ‘Osmocote Exact 3-4 month fertiliser’ (NPK of 21.2:1.9:5.7). Plants were grown under well-watered conditions and variable VPD. Plastic sleeves were used to cover the soil of each pot and minimize soil evaporation. Four pots without plants were used to measure any remaining soil evaporation.

**Fig. 1.**
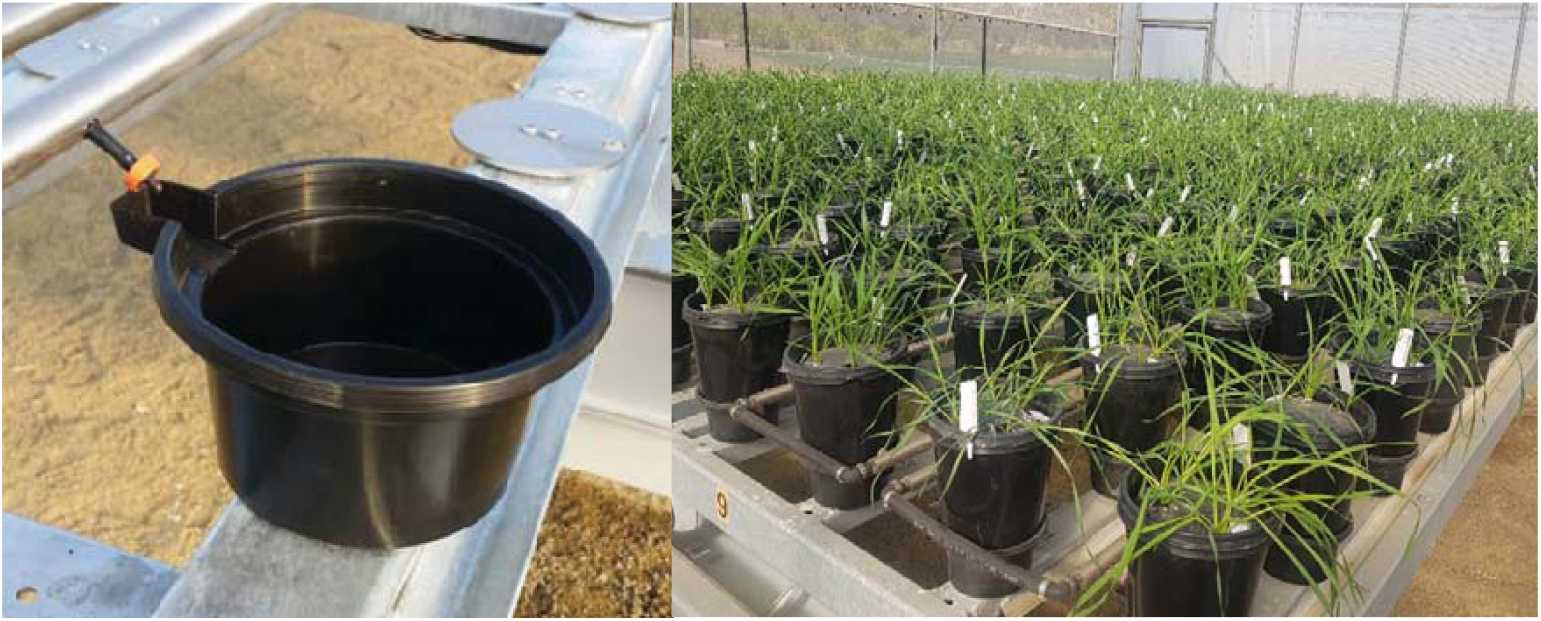
High-throughput lysimeter platform: Water container positioned on a load cell (left) and wheat plants (right).

**Table 1.**
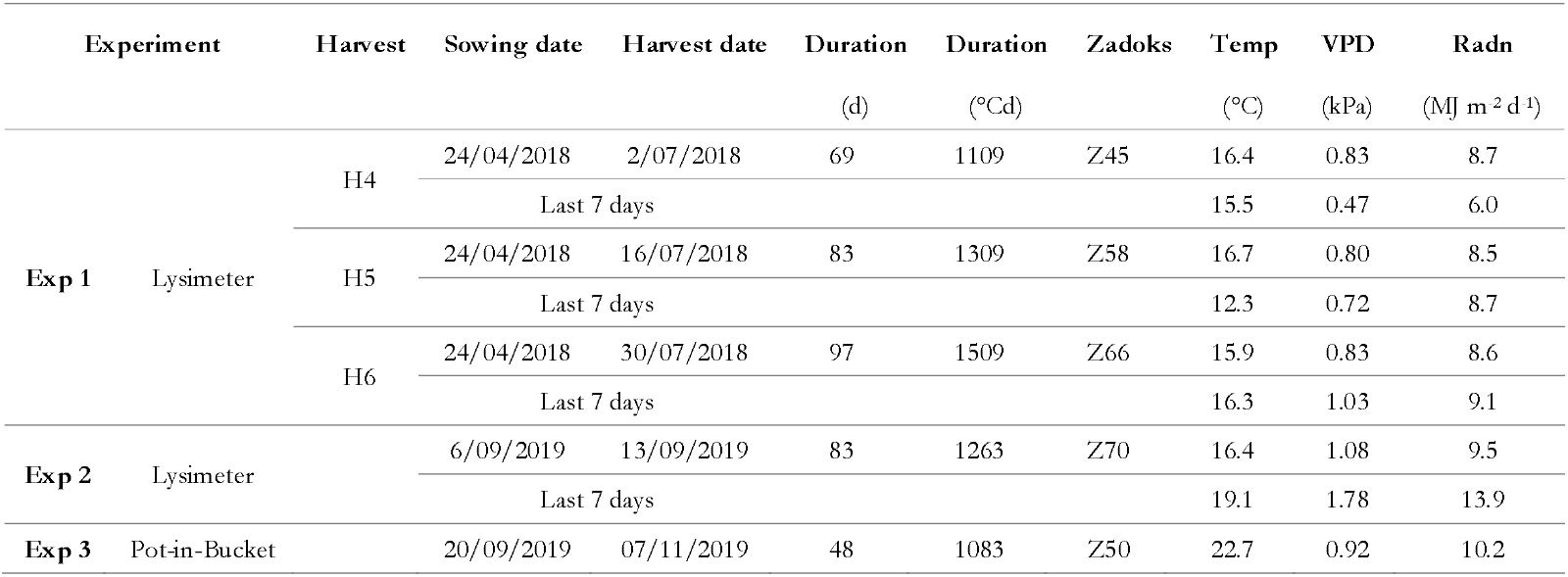
Summary of the three experiments performed in the high-throughput lysimeter platform (Exp 1 in 2018 and Exp 2 in 2019) and in a control environment facility with the Pot-in-Bucket system (Exp 3, 2019). The plant development stage at harvest is reported as the average Zadoks score (Zadoks et al., 1974) across all genotypes. ‘Temp’ is 24-hour average air temperature, ‘VPD’ is 24-hour average air vapour pressure deficit, and ‘Radn’ is 24-hour total solar radiation, averaged from sowing to harvest.

In Exp 1 (2018), six wheat cultivars with contrasting TE (Drysdale, Janz, Mace, Scout, Seri M82 and Suntop) were planted on April 24 in a randomised block design with 6 replications. Plants were harvested six times every fortnight from 200 to 1500^°^Cd after plant emergence (Table 1). In 2019 (Exp 2), 105 wheat genotypes were planted on June 22. These included five groups of genotypes (Supplementary Table S1):

i. seven ‘Reference’ cultivars, namely the six cultivars from Exp 1 and the frequently studied and well-adapted cultivar Hartog.
ii. 74 agronomically-sound spring wheat genotypes that were selected from a ‘Core Collection’ of 372 accessions (which included 120 spring lines) representative of worldwide bread wheat diversity (Balfourier et al., 2007).
iii. four sister lines contrasting for TE (Fletcher, 2020) derived from the Drysdale × Suntop family (hereafter, ‘DS’ lines) of a nested association mapping (NAM) population (Richard, 2015). Both parental lines, Drysdale and Suntop have high TE (Condon et al., 2004; Rebetzke et al., 2009; Tausz-Posch et al., 2012; Fletcher et al., 2018; Collins et al., 2021).
iv. four pairs of near isogenic lines (NILs) varying for a QTL on chromosome 3B involved in yield improvement in hot and dry climates. Selected NIL lines had higher vigour and water use efficiency early in development along with increased biomass, grain number and grain weight following heat stress (Thomelin et al., 2019).
v. 12 advanced drought-adapted ‘CIMMYT’ lines (Reynolds M, personal communication).

All plants were harvested together soon after flowering in the final harvest (H6) of Exp 1 (average genotype at Z66; Zadoks et al., 1974), and when the average genotype was at the beginning of grain development (Z70) in Exp 2.

### 2.2 Experiment with low evaporative demand

In Exp 3 (2019), six wheat cultivars (Drysdale, Janz, Mace, Scout, Seri M82 and Suntop) were grown in a ‘Pot in Bucket’ system (Fletcher et al., 2018) under well-watered and low-VPD conditions in a growth room. The plants, arranged in a randomised block design with six replications, grew in a soil constantly at field capacity. Water losses were recorded weekly. Four seeds of the same genotype were planted 3 cm deep on September 20 in 1.4L ANOVApots® (Anova Solutions) filled with the same potting media as in Exp 1 and 2. Plant emergence occurred on September 25. Seven days after emergence, plants were thinned to keep only two plants per pot so that they grew in a field-like density of 100 plant m^-2^. The same day, a 2-cm layer of white polyurethane beads was added in each pot to reduce moisture loss through evaporation as well as inhibiting weed growth. Four pots were kept without plants to record any remaining water loss. A weather station was used to monitor radiation, relative humidity and air temperature at 10-min intervals. All plants were harvested at heading (Z50, on average) on November 07, 2019.

### 2.3 Plant measurements

Maximum photosynthetic capacity (A_max_) was measured between 9:00 and 15:00 with LI-COR LI-6400 (Exp 1) and CIRAS-3 (Exp 2) portable photosynthesis systems under cloudless conditions for core collection genotypes (Exp 2) and the 6-7 reference cultivars (in Exp 1-2). Measurements were done between July 24 (Z22, on average) and September 12 (Z55, on average) on the penultimate healthy ligulated leaf at about a third of the leaf length (towards the leaf base). Light intensity was set at 1500 µmol m^-1^ s^-1^. Readings were recorded one minute after closing the cuvette head. Additional measurements of stomatal conductance were taken in July 2018 with a porometer (Model SC-1, Decagon Devices Inc., Pullman, WA, USA) with readings taken on 5 days from July 8 to July 16, 2018 between 9:00 and 15:00 on the adaxial surface of the second earliest ligulated healthy leaf at about a third of the leaf length (towards the leaf base). Similar measurements of stomatal conductance were carried out in 2019 (Exp 2) one day before harvest. At each harvest (all Exp), plants were cut at the soil level and green leaf blades, senesced blades, stems with leaf sheaths, and spikes were separated and dried at 70°C for 72 hours before their dry biomass was recorded. Leaf area of the green blades was measured using a LI-3100C leaf area meter (LI-COR, Inc., Lincoln, Nebraska USA).

### 2.4 Data analysis

Shoot TE was calculated as the ratio of dry above-ground dry biomass per gram of water transpired from sowing until harvest in the automated lysimeter (Exp 1-2), and from seven days after emergence (i.e. when the polyurethane beads were added) to harvest in the Pot-in-Bucket experiment (Exp 3).

In Exp 1-2, cumulative water transpired was calculated as the sum of the water loss recorded by weight every 10 min with the lysimeter, minus the irrigation, minus water loss from pots with no plants. Note that, as in most studies on TE, shoot TE rather whole-plant TE was considered as the two have been shown to be generally highly correlated in wheat (Chenu et al., 2018; Fletcher et al., 2018) and sorghum (Chenu et al., 2018; Geetika et al., 2019). Environmental effects on transpiration rate were analysed in detail by using 10-min-interval VPD and transpiration rates in Exp 1 and Exp 2. In order to account for genotypic variation over time in the effect of canopy size on transpiration rates, a novel approach was taken in which transpiration rates were normalised each day, for each pot, with the transpiration rate corresponding to a reference VPD of 1.2 kPa. This transpiration rate was estimated with a linear interpolation between transpiration rates measured every 10 min between 8:00 and 11:00 am. To reduce the impact from the limited sensitivity of the lysimetric platform, the analysis was performed on data from 750°Cd after sowing onwards, i.e. not considering data for small canopy size in the early growth stage. In addition, for each pot, cumulative transpiration for the last 7-d of the experiment (pre-harvest) was normalised by the green leaf area (GLA) measured at harvest and cumulated for every 10-min time step when VPD≥1.3 kPa and Radn≥0.84 MJ m^-2^ h^-1^ (TR7_GLAnorm-highVPD_).

Data analysis and presentation were carried out with the R programming language (R Core Team, 2023) and the R package ‘ggplot2’ (Wickham, 2016). Analyses of variance (ANOVA) were performed to assess the effects of genotype, experiment, harvest, and their interactions on measured traits. Pearson and Spearman correlation coefficients were calculated to examine relationships among traits and environmental variables across experiments and genotype groups.

Where relevant, linear regression analyses were performed to compare slopes and intercepts of trait–environment relationships between genotypes or genotype groups. Differences among genotype means were considered significant at P<0.05 unless otherwise stated. In Exp 2, the ‘k-means’ clustering algorithm implemented in the R package ‘stats’ was used to classify genotypes based on their average patterns of hourly normalised transpiration rates.

## 3 RESULTS

### 3.1 Genotype ranking for transpiration efficiency maintained across environments

Genotypic variations for TE were significant (P<0.001) in both Exp 1 and Exp 2 (Fig. 2). In an analysis of variance, *Experiment* and *Harvest* also had effects on TE (P<0.001), but *Experiment × Genotype* and *Harvest × Genotype* interactions were both statistically insignificant (P>0.10). Harvest before 900_o_Cd after sowing (i.e. Exp 1 H1-H3) presented greater intra-genotypic variability for TE than later harvests likely due to the relatively smaller size of plants and the level of resolution from the lysimeter. Exp 1-H5 recorded a higher average TE (5.3 g kg^-1^) than Exp 1-H4 (4.6 g kg^-1^) and Exp 1-H6 (4.8 g kg^-1^; Fig. 2). Exp 2 was performed with a later sowing date than Exp 1 under higher VPD and radiation and had a slightly higher average TE (5.1 g kg^-1^) than in Exp 1 for a harvest at similar development stage (4.8 g kg^-1^ in Exp 1-H6).

**Fig. 2.**
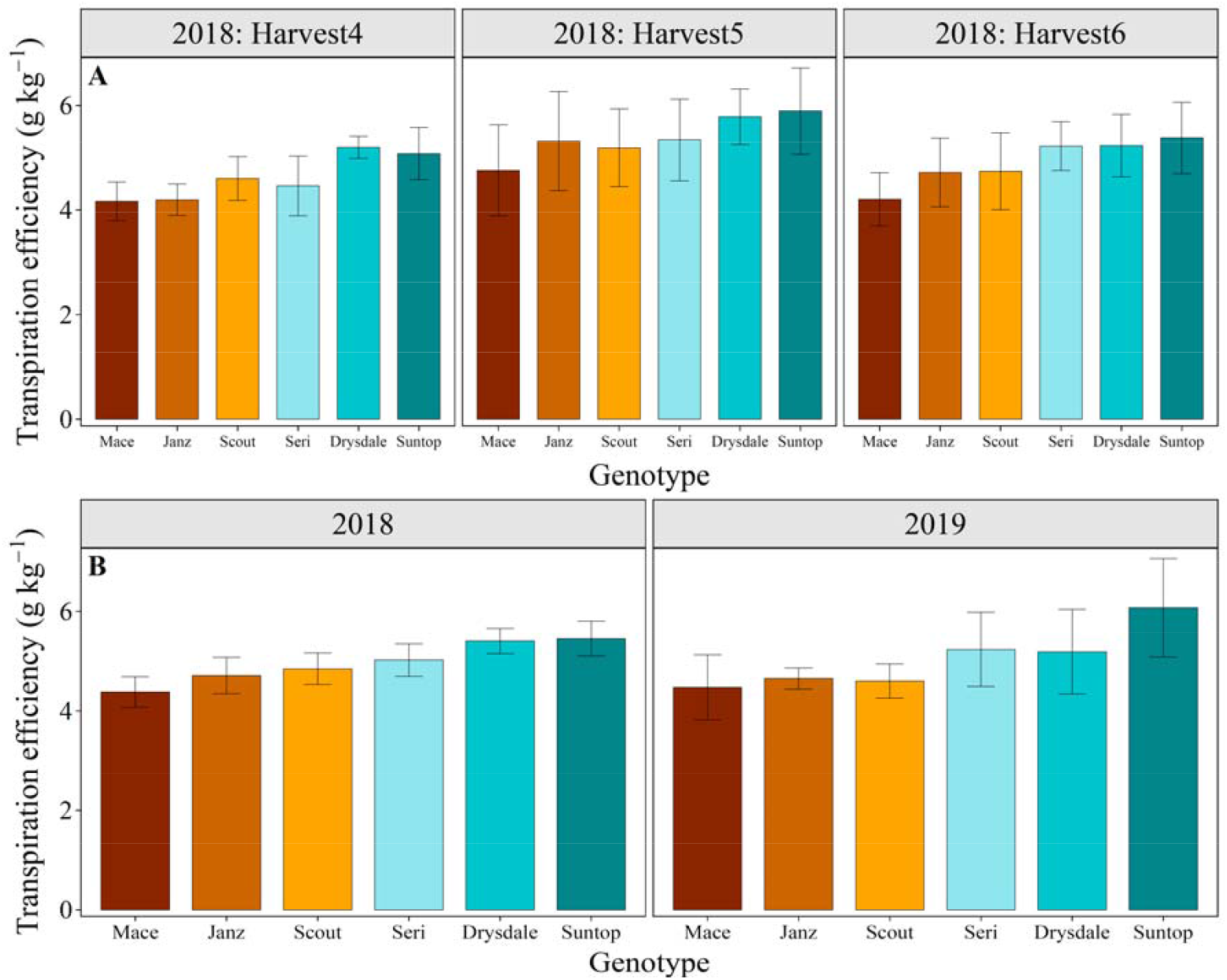
Transpiration efficiency (TE) of six reference genotypes in 2018 (Exp 1) and 2019 (Exp 2) with (A) TE for harvests 4, 5 and 6 (i.e. harvests occurring after 900^°^Cd after sowing) in 2018, and (B) average TE across these three harvests in 2018 and (C) TE for the post-flowering harvest from 2019. Genotypes are sorted based on average TE in 2018. Error bars correspond to 95% confidence intervals.

High correlations for TE among genotypes were observed across environments (Fig. 2). TE values from the three latest harvests in Exp 1 were significantly correlated with each other (r=0.86, on average) and with Exp 2 (r=0.81 and spearman rank correlation of 0.89 with the average TEs across Exp 1 H4-H6). Overall, Mace had consistently the lowest TE (4.2 g kg^-1^ in both Exp 1 and Exp 2), followed by Janz, Scout and Seri, while Drysdale and Suntop were consistently the highest TE lines (Fig. 2). TE thus ranged from 4.2 g kg^-1^ in both Exp 1 and Exp 2 (Mace) to 5.3 g kg^-1^ in Exp 1 and 6.0 g kg^-1^ in Exp 2 (Suntop; Fig. 2B).

### 3.2 Transpiration efficiency and VPD response of transpiration rate

Component traits of transpiration efficiency include traits related to biomass accumulation, such as photosynthesis, and traits related to transpiration, such as stomatal conductance. Limited genotypic variations were observed for maximum photosynthetic capacity (A_max_) in the six studied genotypes. In Exp 1, Mace tended to have a lower A_max_ compared to the other genotypes (Fig. 3A), while in Exp 2, it had a photosynthetic capacity similar to the other genotypes (Fig. S21).

**Fig. 3.**
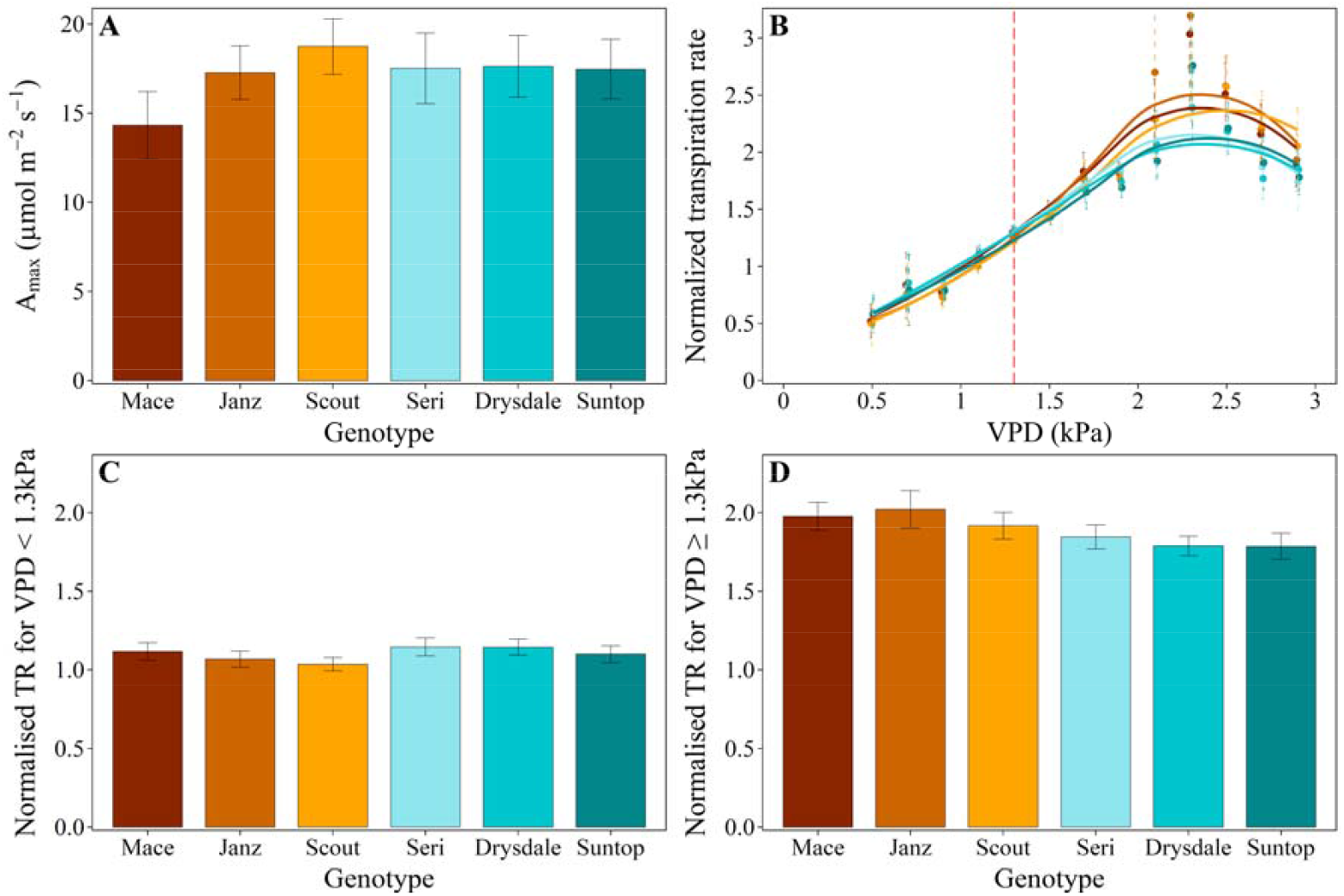
Physiological traits associated with transpiration efficiency for six reference genotypes in 2018 (Exp 1): (A) maximum photosynthetic capacity (Amax), (B) normalised transpiration rate (TR_norm_) in response to VPD, (C) average normalised transpiration rate at low VPD (<1.3 kPa; TR_norm-lowVPD_), and (D) average normalised transpiration rate at high VPD (≥1.3 kPa; TR_norm-highVPD_). The dot red vertical bar in (B)corresponds to a VPD of 1.3 kPa. Error bars correspond to 95% confidence intervals. Data for Exp 2 (2019) are presented in Supplementary Fig. S2.

Limited genotypic variations were observed for stomatal conductance at the tissue level (Fig. 4). Under radiation-limited conditions, stomatal conductance was strongly related to radiation (Fig. 4A). No significant variation was observed in this response among the six studied genotypes (P>0.05). Under higher radiation (≥0.84 MJ m^2^ h^-1^, typically between 9:00 and 15:00 on a sunny day), stomatal conductance increased linearly with VPD for all six genotypes up to a breakpoint around a VPD of 1.3 kPa (from 1.27 to 1.32 kPa depending on the genotype), and then levelled off at 350-400 mmol m^2^ s^-1^ (Fig. 4B). All six genotypes responded in a similar way and no significant differences were observed in this response.

**Fig. 4.**
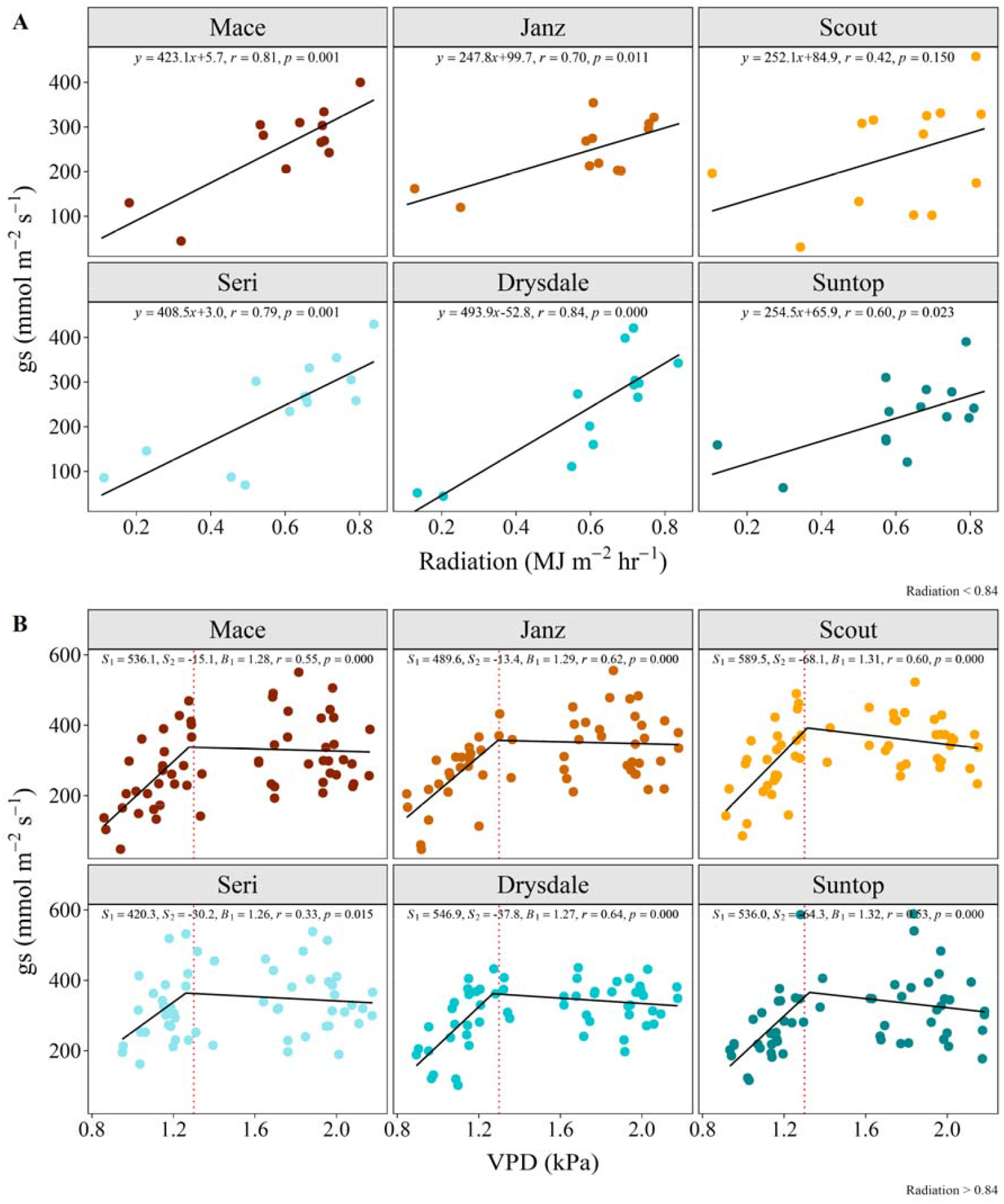
Stomatal conductance (gs) against (A) radiation intensity in low-radiative conditions (Radn<0.84 MJ m^-2^ *h*^-1^) and against (B) VPD in high-radiative conditions (Radn≥0.84 MJ m^-2^ h^-1^) for the six reference genotypes. Measurements of gs were taken with porometer between 9:00 and 15:00 on six sunny days in July 2018 (Exp 1). The dot red vertical bar in (B) corresponds to a VPD of 1.3 kPa.

By contrast, significant genotypic variations were observed when analysing the normalised transpiration rate (TR_norm_) at the plant level. Sub-daily values of transpiration rates were normalised with a daily reference transpiration rate for a VPD of 1.2 kPa, for which (and below which) no genotypic variability was observed. This normalisation enabled accounting for variation in canopy size across genotypes and over different growth stages. In addition, based on the observed response of stomatal conductance to low radiation (Fig. 4A), the analysis focused on high-radiation conditions (≥0.84 MJ m^2^ h^-1^). Genotypic differences were observed for VPD≥1.3 kPa (Fig. 3B, D; Supplementary S1B, D), a threshold that also corresponded to the VPD break point observed for stomatal conductance at the tissue level (Fig. 4B). Under high VPD conditions, Mace, Janz and Scout had consistently higher TR_norm_ values than Seri, Drysdale and Suntop (Fig. 3B, D).

In both experiments for high radiation levels (Radn≥0.84 MJ m^2^ h^-1^), significant genotypic differences were found in the average normalised transpiration rate for VPD≥1.3 kPa (TR_norm-highVPD_), while no substantial variations were found for normalised transpiration rate for VPD<1.3 kPa (TR_norm-lowVPD_; Fig. 3; Supplementary Fig. S1). The average normalised transpiration for VPD≥1.3 kPa (TR_norm-highVPD_) of Mace, Janz and Scout was higher than those of Seri, Drysdale and Suntop (Fig. 3D) and significant and high genotypic correlations for TR_norm-highVPD_ were observed between Exp 1 and Exp 2 (r=0.82; Fig. 3D; Supplementary Fig. S1). Overall, genotypes with higher TR_norm-highVPD_ tended to have lower TE, while genotypes with low TR_norm-highVPD_ tended to have higher TE (Fig. 2 and Fig. 3B-D). Average TR_norm-highVPD_ explained 46% of TE variations in Exp 1 and 17% in Exp 2 for the six reference genotypes.

For high VPD and radiation levels (VPD≥1.3 kPa and Radn≥0.84 MJ m^2^ h^-1^), normalised transpiration rate was significantly correlated to the cumulative transpiration over the last 7 days of the season normalised by green leaf area at harvest (TR7_GLAnorm-highVPD_; g mm^-2^) in both Exp 1 (r=0.57, P=0.014) and Exp 2 (r=0.43, P=0.015 for the elite lines; Supplementary Fig. S2B).

At low VPD (Exp 3), relatively limited genotypic variability were observed for TE given high intra-genotypic variations (Fig. 5A). Drysdale and Suntop tended to have greater intrinsic TE than the other genotypes, and had a TE significantly higher than Scout. By contrast to the other experiments, Suntop had a relatively low above-ground dry biomass, cumulative transpiration, and green leaf area in this experiment (Fig. 5B-D).

**Fig. 5.**
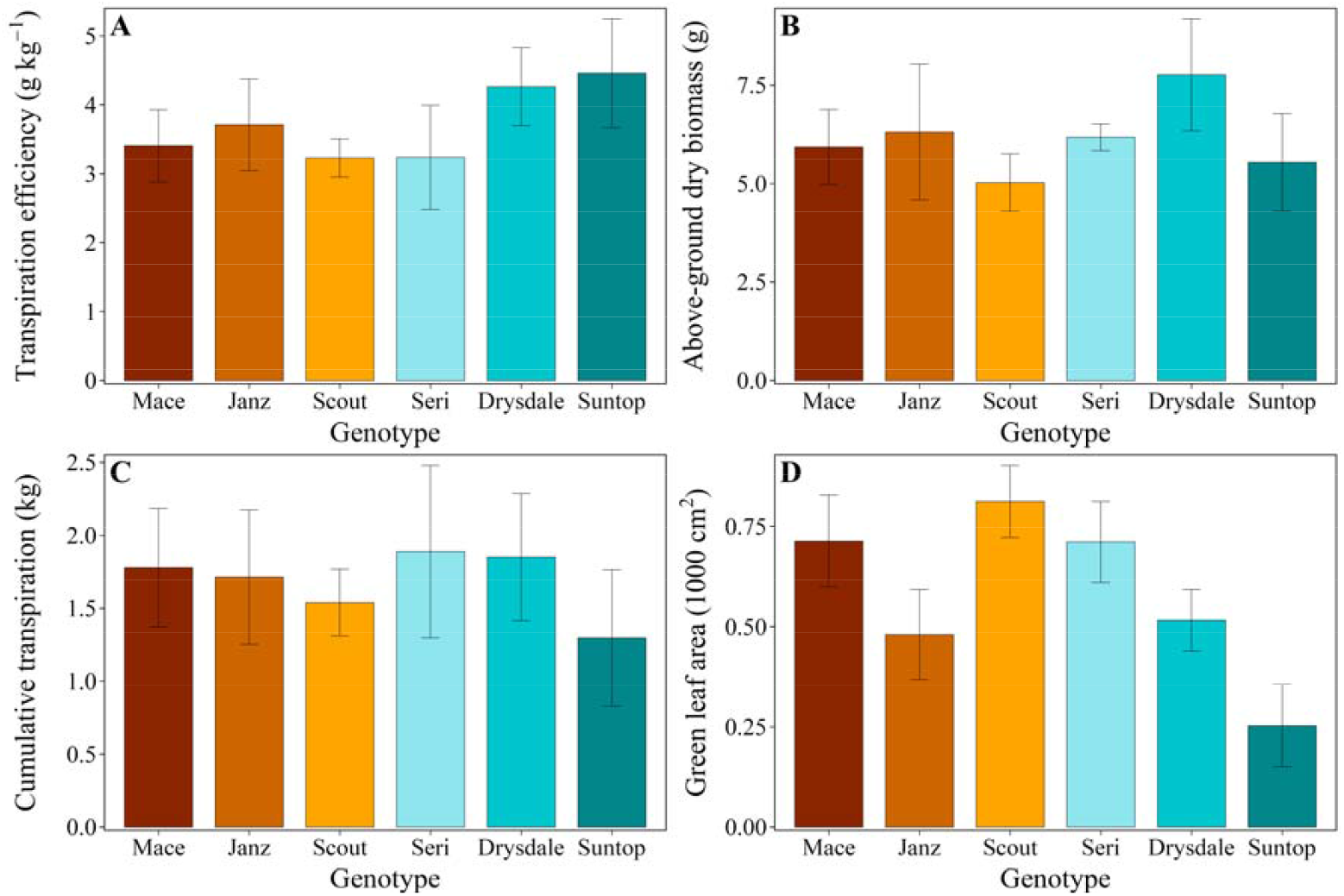
Intrinsic transpiration efficiency and associated traits for six cultivars grown under low VPD conditions: (A) transpiration efficiency, (B) above-ground dry biomass per plant, (C) cumulative transpiration and (D) green leaf area in Exp 3. Error bars correspond to 95% confidence intervals.

### 3.3 Genetic variations in transpiration efficiency, photosynthetic capacity and normalised transpiration rate

A broad range of genotypes including lines from a core-collection, elite drought-tolerant CIMMYT lines, contrasting-TE sister lines derived from crosses between high-TE parents Suntop and Drysdale (‘SD lines’), near-isogenic lines for a QTL involved in early vigour and yield increase under heat (‘NIL’) and benchmarking cultivars (‘Reference’) were phenotyped in Exp 2. Genetic variations in TE ranged from extremely low (2.91 g kg^-1^) to extremely high (7.74 g kg^-1^) across the 105 lines (Fig. 6A,Fig. 7). In an analysis of variance, significant genotype and genotype-group effects were found for all the studied traits, including TE, biomass accumulation, cumulative transpiration, and maximum photosynthesis capacity (P<0.001). DS and Core Collection lines had the highest average TEs among the studied groups of genotypes (6.21 and 5.30 g kg^-1^, respectively), followed by NIL (5.13 g kg^-1^), Reference (5.03 g kg^-1^) and CIMMYT (4.92 g kg^-1^) genotypes. Large variations in TE and its component traits were observed among genotype groups.

**Fig. 6.**
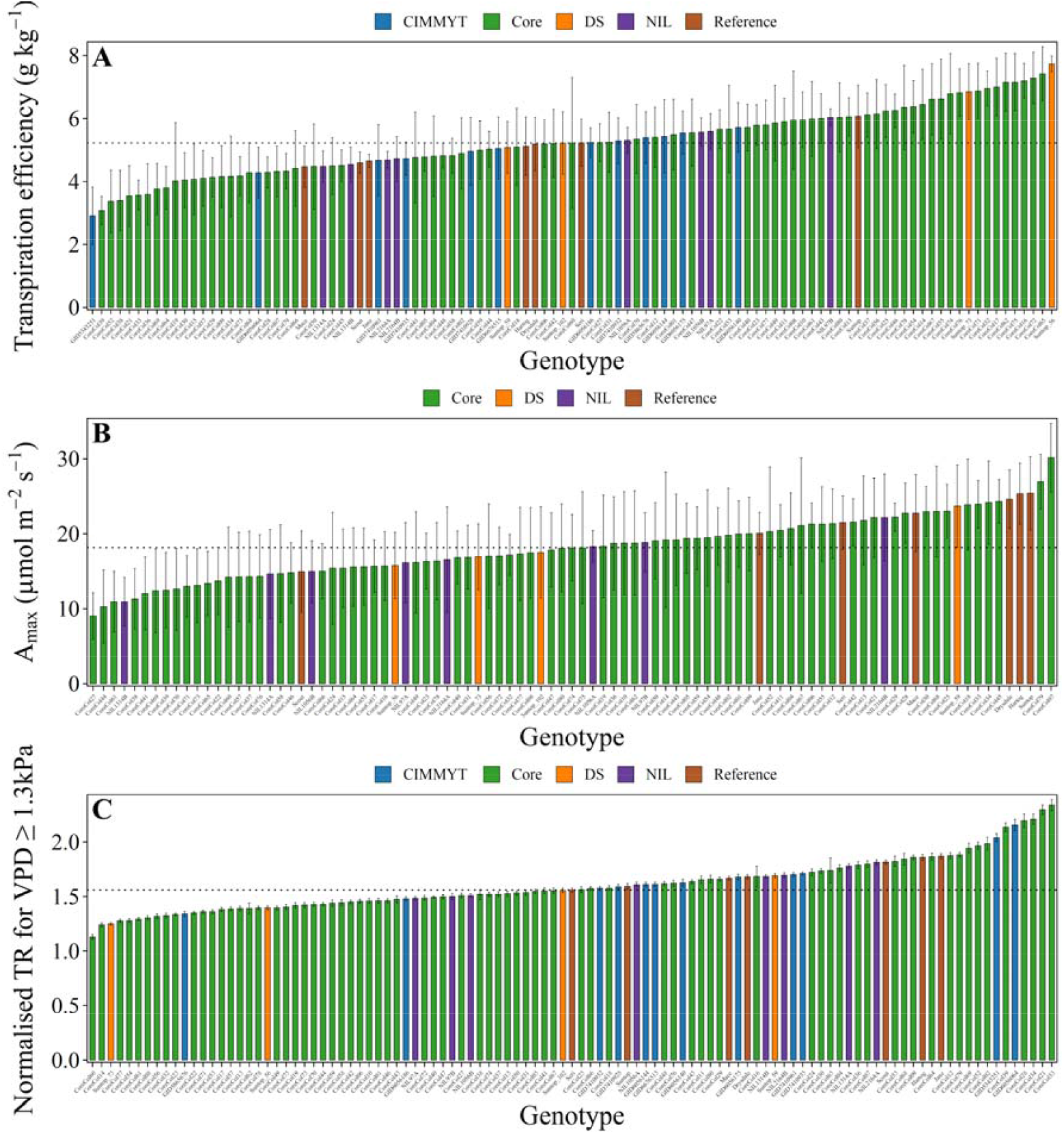
Physiological traits associated with transpiration efficiency across 105 genotypes: (A) Transpiration efficiency, (B) maximum photosynthetic capacity (Amax) and (C) average normalised transpiration rate (TR) under high evaporative demand in Exp 2. Normalised transpiration rate for VPD≥1.3 kPa and Radn≥0.84 MJ m^-2^ h^-1^ were average between 750^°^Cd after sowing to harvest. In each panel, the dotted line corresponds to the median value. Error bars correspond to 95% confidence intervals. Abbreviations: CYMMYT, advanced drought adapted lines from CYMMYT; Core, core collection lines representative of worldwide bread wheat diversity; DS, four sister lines from the Drysdale × Suntop family; NIL, four pairs of near isogenic lines varying for a QTL on chromosome 3B; and Ref, seven ‘reference’ Australian cultivars.

**Fig. 7.**
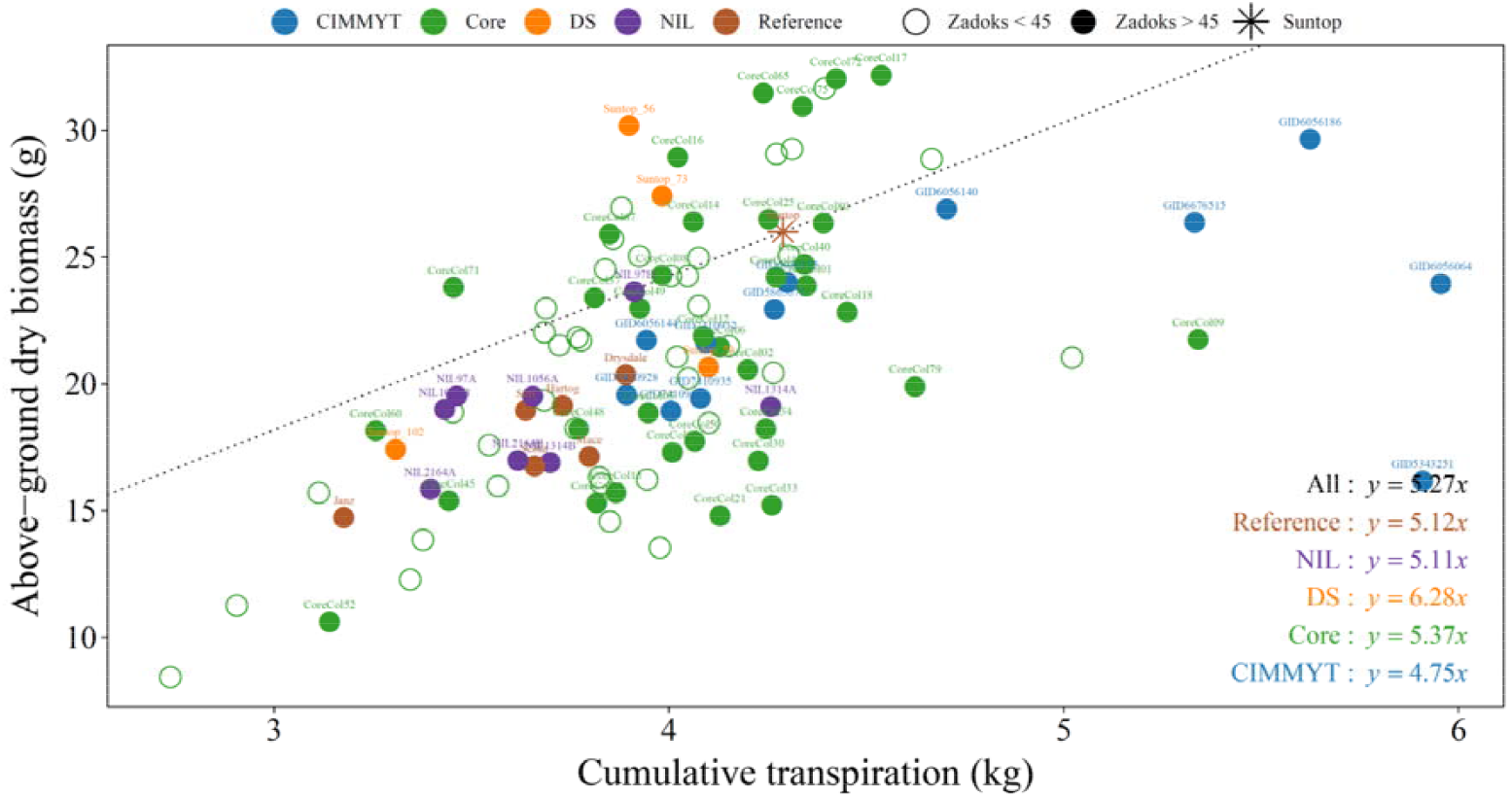
Relationship between cumulative transpiration and above-ground dry biomass per plant at harvest for 105 genotypes in Exp 2. Open symbols correspond to genotypes with a slow phenology (not flowering at harvest), while filled symbols correspond to lines with a phenology similar to the reference cultivars. The slope of the dashed line corresponds to the TE of high-TE benchmark cultivar Suntop. Points above this line correspond to genotypes with higher TE than Suntop.

For TE, the two high-TE DS lines (Suntop_56, 7.74 g kg^-1^; and Suntop_73, 6.85 g kg^-1^) were among the genotypes of the highest TE values (Fig. 6A), well above their parents Drysdale (5.19 g kg^-1^) and Suntop (6.07 g kg^-1^). Among the top 10 genotypes were also CoreCol65 (7.42 g kg^-1^), CoreCol72 (7.29), CoreCol16 (7.20), CoreCol75 (7.16), CoreCol62 (7.15), CoreCol17 (7.00), CoreCol32 (6.96), and CoreCol71 (6.88 g kg^-1^).

Maximum photosynthesis capacity (A_max_) ranged from 9.0 to 30.2 µmol m^-2^ s^-1^ in the lines for which A_max_ was measured (i.e. all except CIMMYT lines), with three reference cultivars and one DS line among the top 10 (Fig. 6B). A_max_ was not correlated with TE (P>0.20, Fig. 8A).

**Fig. 8.**
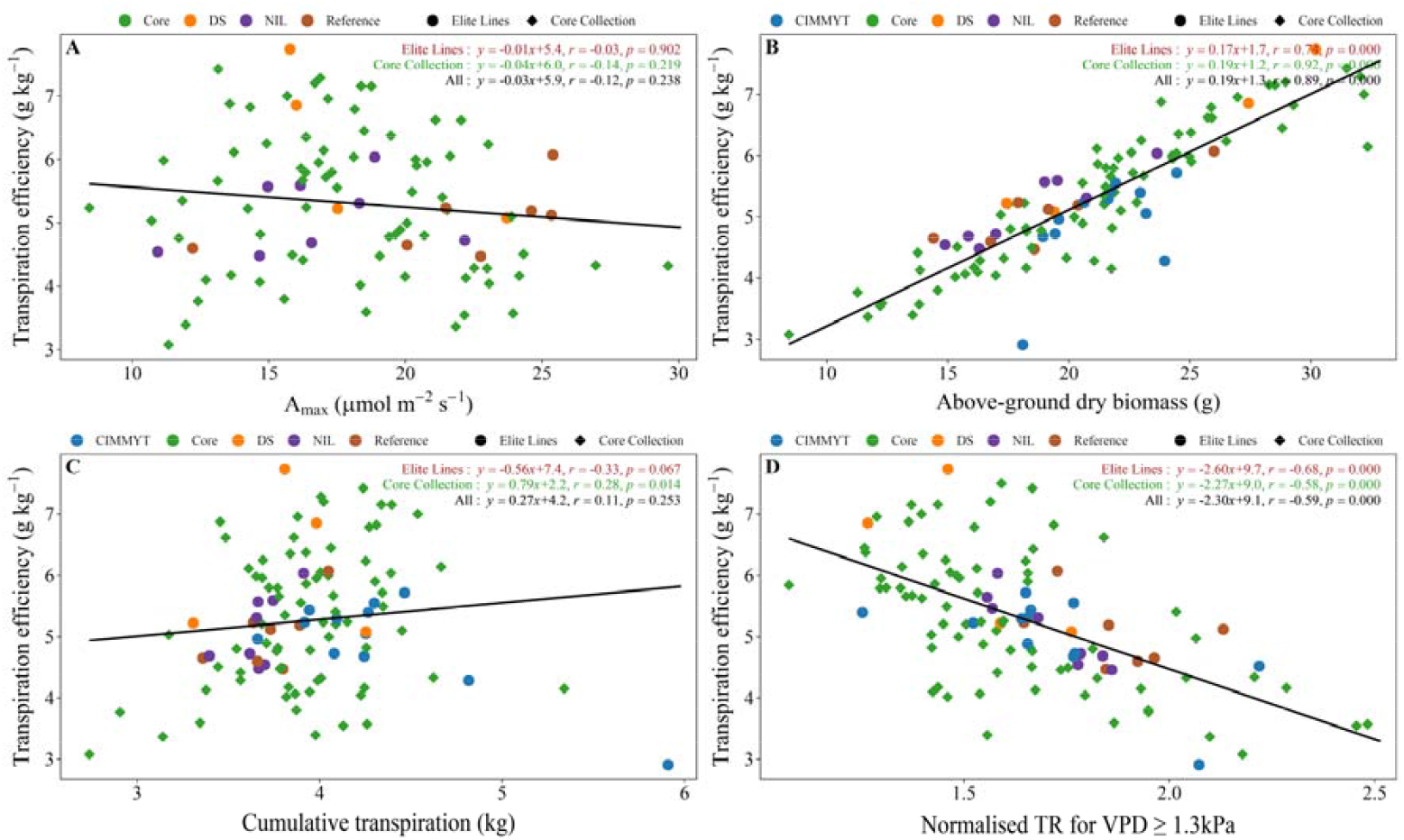
Correlations between transpiration efficiency and (A) maximum photosynthetic capacity (A_max_), (B) above-ground dry biomass per plant, (C) cumulative transpiration, or (D) normalised transpiration rate (TR) for VPD≥1.3 kPa and Radn≥ 0.84 MJ m^-2^ *h*^-1^ (TR_norm-highVPD_) across studied genotypes in Exp 2. ‘Elite Lines’ include elite breeding lines from CIMMYT lines, NIL and DS lines (which have Australian cultivars as genetic backgrounds) and the reference cultivars.

Above-ground dry biomass ranged from 8.4 to 32.3 g plant^-1^ in the tested lines, with two DS lines (Suntop_56 and Suntop_73) among the top lines. Above-ground dry biomass per plant was strongly correlated with TE (r=0.48, P<0.05 in Exp 1; r=0.89, P<0.001 in Exp 2, Fig. 8B).

Cumulative transpiration ranged from 2.74 to 5.91 kg plant^-1^ in the 105 tested lines, with one DS line (Suntop_102), one reference cultivar (Janz) and one NIL line (NIL2164A) along with seven Core Collection lines among the top 10 lines with minimum TR_norm-highVPD_. While cumulative transpiration was not significantly associated with TE (P>0.01, Fig. 8C), the correlation between average TR_norm-highVPD_ and TE was strong (r=-0.59, P<0.0001, Fig. 8D). TE was also correlated significantly with TR7_GLAnorm-highVPD_ (r=-0.28, P=0.004; Fig. S3B), as TR_norm-highVPD_ was relatively strongly correlated with TR7_GLAnorm-highVPD_ (r=0.43 and P=0.015 for Elite lines; r=0.41 and P<0.001 for Core Collection lines; Supplementary Fig. S3A), and with stomatal conductance (r=0.28, P<0.001, tested across the Reference cultivars and DS genotypes). For the 105 genotypes, TR_norm-highVPD_ and TR7_GLAnorm-highVPD_ explained 35 and 8% of the variations in TE, respectively.

### 3.4 High TE lines tend to have low TR_norm-highVPD_

The 10 genotypes with the highest, median and lowest TE values tended to have low, median and high value of TR_norm-highVPD_, respectively (Fig. 9A). While these groups had average TE of 7.26, 5.23 and 3.72 g kg^-1^, respectively (Fig. 9C), they had an average TR_norm-highVPD_ of 1.41, 1.48 and 1.84, respectively (Fig. 9E), with the high-TE group having significantly lower TR_norm-highVPD_ than the other groups.

**Fig. 9.**
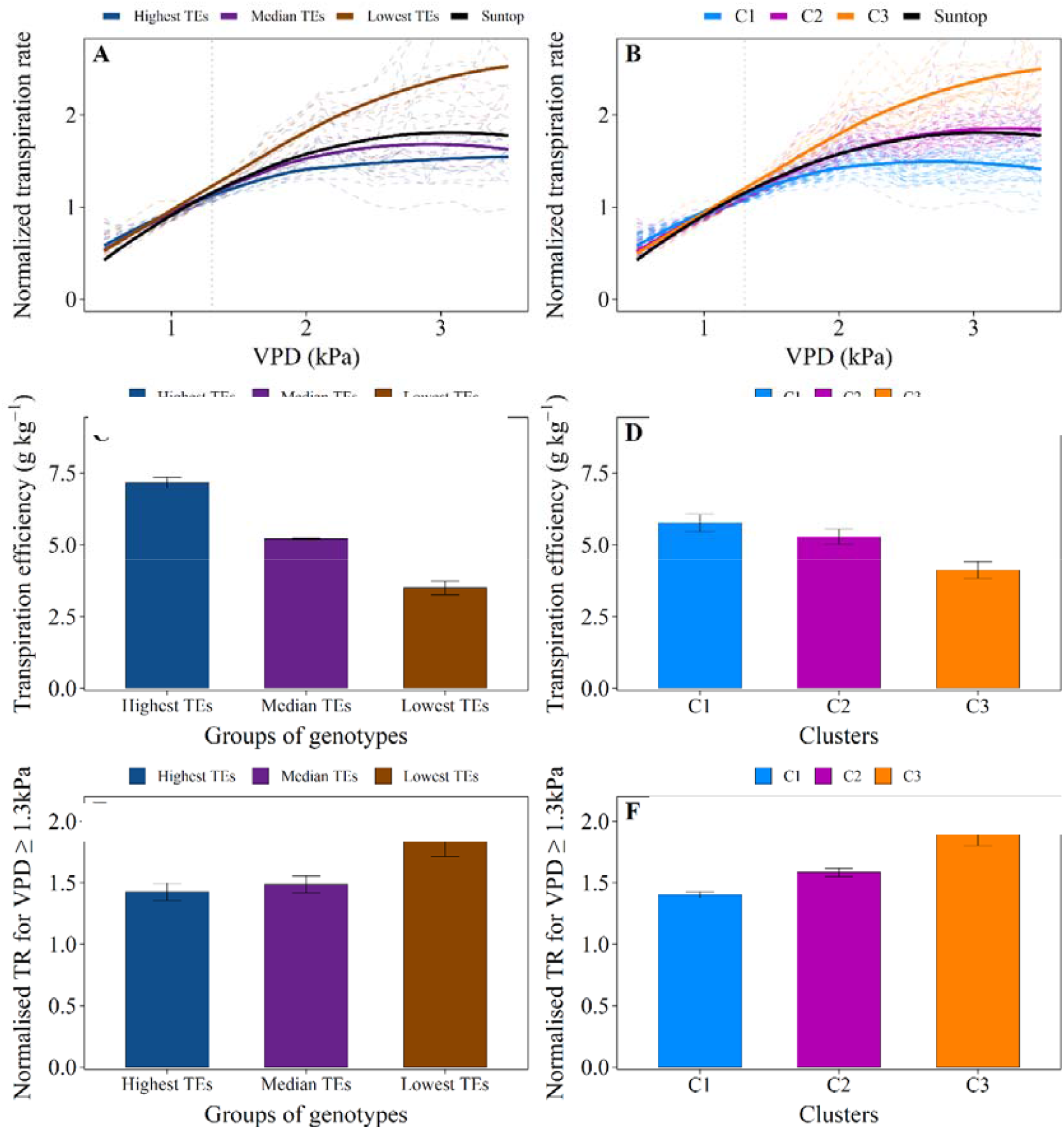
(A-B) Normalised transpiration rate (TR_norm_) response to VPD, (C-D) transpiration efficiency (TE) and (E-F) normalised transpiration rate (TR) at high evaporative demands (TR_norm-highVPD_) for genotypes with contrasting TE (A, C, E) and three clusters of genotypes contrasting in TR_norm-highVPD_ (B, D, F) in Exp 2. In (A, C, E), groups of genotypes with contrasting TE correspond to the 10 genotypes with the highest, median and lowest TE averages. In (B, D, F), the 105 genotypes were clustered in groups (‘clusters’) based on responses of normalised transpiration rate to VPD. In (E-F), normalised transpiration rate are presented for VPD≥1.3 kPa and Radn≥0.84 MJ m^-2^ h^-1^ averaged between 750^°^Cd after sowing and harvest. Error bars correspond to 95% confidence intervals.

Genotypes were also clustered based on their responses of normalised transpiration rate to VPD (Fig. 9B). The three resulting clusters (C1, C2, C3) varied in TR_norm-highVPD_, with (i) the low TR_norm-highVPD_ cluster (C1; largest decrease in TR_norm_ when VPD≥1.3 kPa compared to C3) had the greatest average TE (5.9 g kg_-1_), with most of its genotypes having a well-below-median TR_norm-highVPD_ and a well-above-median TE (Supplementary **Fig. S4**), (ii) the medium TR_norm-highVPD_ cluster (C2; median decrease in TR_norm_ when VPD≥1.3 kPa compared to C3) had an average TE of 5.1 g kg^-1^, and (iii) the high TR_norm-highVPD_ cluster (C3; near linear TR_norm_-VPD response) had the lowest TE (4.1 g kg^-1^), with most of its genotypes being in the lower tail of the TE ranking (Supplementary **Fig. S4**A) and all being in the top tail of the TR_norm-highVPD_ ranking (Supplementary **Fig. S4**B).

In order to further investigate environmental effects on TE, the TE-TR_norm_ relationship was analysed over various ranges of VPD (0-1.3, 1.3-1.5, 1.5-2.0, 2.0-2.5, and 2.5-3.0 kPa) across all genotypes in Exp 1 and Exp 2 (Fig. 10). To that end, the average TR_norm_ was calculated over selected ranges of VPD. Highly significant correlations were observed for all high VPD ranges (VPD ≥1.3 kPa) but not for VPD<1.3 kPa. This finding further underscores the greater significance of transpiration rate under high rather low VPD conditions. It also suggest that reducing transpiration rates during periods of moderately high sub-hourly VPD may be more critical than during extremely high VPD events, given their larger cumulative contribution to seasonal water use.

**Fig. 10.**
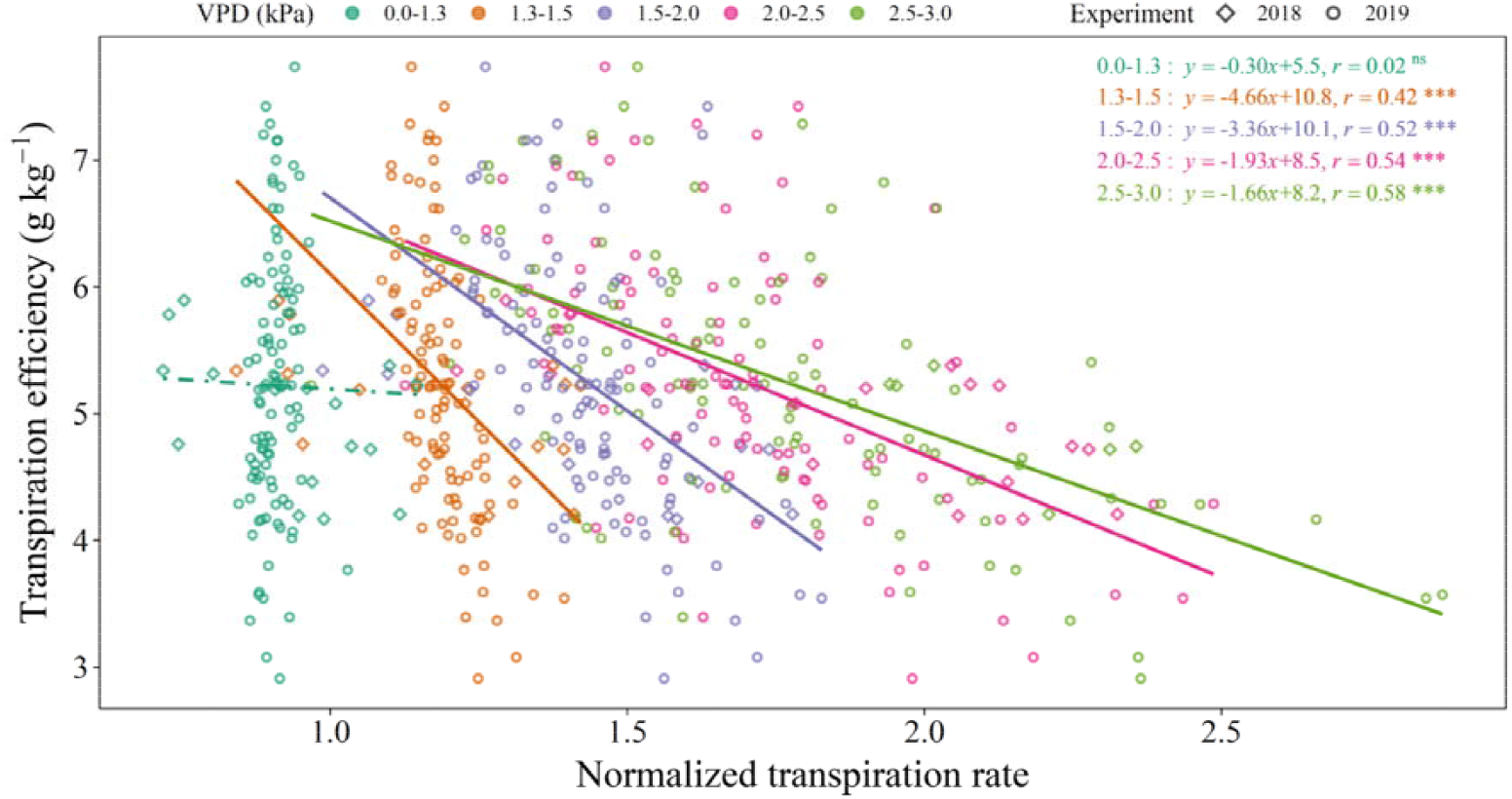
Transpiration efficiency against normalised transpiration rate (TR_norm_) for different evaporative demand ranges. For each genotype, normalised transpiration rates were averaged for different VPD ranges (0-1.3, 1.3-1.5, 1.5-2.0, 2.0-2.5, and 2.5-3.0 kPa) when Radn≥0.84 MJ m^-2^ h^-1^ between 750^°^Cd after sowing and harvest. Data pooled from Exp 1 and Exp 2. ***: P<0.001, **: P<0.01, *: P<0.05, ns: not significant.

## 4 DISCUSSION

### 4.1 Transpiration efficiency (TE) and its components: a high correlation between TE and normalised transpiration rate at high evaporative demand

Defined by the amount of above-ground dry biomass produced per unit of water transpired, plant-level TE was more closely related to above-ground biomass (significant correlation of 0.89; Fig. 8B) than plant cumulative transpiration in a panel of 105 genotypes (and r=0.11 Fig. 8C), revealing similar findings as in sorghum (Xin et al., 2009). At the tissue level, TE is defined as the ratio of CO_2_ assimilation via photosynthesis to water loss due to transpiration (Condon, 2020). Yet, no significant correlation was found between plant-level TE and A_max_ across the 105 studied lines (Fig. 8A). By contrast, TE was strongly correlated with plant-level TR_norm-highVPD_ (r=-0.59; Fig. 8D) and genotypes with the lowest plant-level TR_norm-highVPD_ were those with greatest TE (Fig. 9; Supplementary Fig. S3).

In fluctuating environments, wheat genotypes were found to differ in normalised transpiration rates for VPD≥1.3 kPa in both Exp1 (Fig. 3B) and Exp 2 (Supplementary Fig. S1B), which is much lower than VPD thresholds of 2.4-3.9 kPa reported for different wheat genotypes in controlled environments (Schoppach and Sadok, 2012). Note a VPD of 1.3 kPa corresponds to an air temperature of 26^°^C with 60% relative humidity, or 22^°^C with 50% relative humidity, i.e. conditions that are relatively frequent in wheat production environments (e.g. over 60% of the 10 min data recorded in Exp 1 under high radiation, i.e. between ∼9:00 and ∼15:00). By contrast, a VPD of 2.4 kPa is the equivalent of an air temperature of 32^°^C with 50% relative humidity, or 36^°^C with 60% relative humidity, i.e. conditions rarely occurring in environments where wheat is cultivated. Interestingly, a breakpoint at 1.3 kPa was also observed for the response of stomatal conductance to VPD (Fig. 4B) but no genotypic differences could be detected for this response due to high intra-genotypic variations. This VPD threshold was close to the 1.07 kPa threshold (averaged across 17 lines) reported in durum wheat (Medina et al., 2019).

While only significant between most-contrasting genotypes, differences in TE observed under low VPD (Exp 3; Fig. 5A) suggest that genotype variations in TE are not only driven by response to high VPD. TE was highly correlated to above-ground dry biomass but not photosynthetic capacity (Amax), suggesting that differences in light interception and possibly plant architecture may be involved. In sorghum, a strong relationship was found between plant-level conductance and radiation for low-radiation levels (Geetika et al., 2019). Accordingly, in the current study, stomatal conductance was predominantly controlled by radiation under low-radiation conditions (Radn<0.84 MJ m^-2^ h^-1^, Fig. 4A), while under high-radiation conditions, stomatal conductance was strongly impacted by VPD. Light is known to be one of the most dynamic environmental signals influencing both photosynthetic activity and stomata conductance but at different speed (Lawson and Vialet-Chabrand, 2019). A variation in radiation (e.g. a cloud passing) results in a steady state of photosynthetic rate within several tens of seconds to minutes, whereas changes in stomatal conductance can take minutes to hours (Barradas & Jones, 1996; Lawson & Morison, 2004; Lawson et al., 2010; Vico et al., 2011; McAusland et al., 2016) thus influencing TE in fluctuating environments such as in Exp 1 and 2. Other factors such as light quality and CO_2_ also impact stomatal conductance and transpiration (e.g. Yu et al., 2004; Bourgault et al., 2013; Gawinowski et al., 2025). The impact of such environmental factors, their interactions and genetic variability should be further investigated across scales to better understand crop responses in current and future climates.

### 4.2 A new method to screen limited transpiration efficiency at high VPD in environments with fluctuating evaporation demand

TE was screened robustly, with consistent genotype ranking across environments for growth stages beyond 900^°^Cd after sowing (r=0.89 between Exp 1 and Exp 2, Fig. 1) consistently with the findings from Fletcher et al. (2018). TR_norm-highVPD_ was highly correlated to TE across environments (r=-0.68 in Exp 1 and r=-0.59 in Exp 2) and can be considered as a surrogate trait for TE. Importantly, phenotyping of TR_norm-highVPD_ also provided consistent genotype ranking (r=0.82 between Exp 1 and Exp 2) and was easier to measure than TE, with measurements taken automatically by the lysimeter platform and no requirement for biomass sampling or estimation.

Other methods exist to screen TE with automatic lysimeter platforms (e.g. Vadez et al., 2015; Ryan et al., 2016; Chenu et al., 2018) or the low-cost, low-technology Pot-in-Bucket system (Fletcher et al., 2018) particularly relevant for developing countries, but all require additional resources to estimate biomass (to calculate TE) and/or green leaf area (to normalise transpiration rate by green leaf area (GLA); Vadez et al., 2015). Considerable processes in high-throughput imaging technologies have been made to estimate plant leaf area (Fanourakis et al., 2014; Vadez et al., 2015; Prado et al., 2018) yet precise phenotyping of genetic differences remains difficult in species like wheat due to high tillering and leaf overlapping. In addition, recent research has revealed the importance of plant density and radiation on TE (Pillioni et al., 2024), which poses issue for 3D imaging systems where pots need to be isolated to capture genotype architectural characteristics and GLA (e.g. Xue et al., 2024). By contrast, in the proposed screening method, normalising transpiration rate based on a reference value (i.e. pot weight at low VPD) updated daily allows the capture of canopy-size variation and enables TR_norm-highVPD_ estimation in a system where plants can be grown in densities similar to production systems, with leaves overlapping from different genotypes (i.e. between different pots).

The proposed method was used under fluctuating environments with radiation, temperature and VPD dynamics similar to production environments. This contrasts from phenotyping done in controlled environments for VPD levels maintained over time (e.g Schoppach and Sadok, 2012). The framework allowed genotype discrimination for relatively low VPD thresholds (1.3 kPa in Fig. 3B and Supplementary Fig. S1 vs. 2.4-3.9 kPa in Schoppach and Sadok, 2012). Given the time differences required for photosynthesis (several seconds to minutes) and stomatal conductance (several minutes to hours) to adjust to rapid change in radiation (Lawon and Vialet-Chabrand, 2019), working in natural environments mimicking production systems appears to have greater relevance for breeding purposes.

High-throughput estimations of TE have also been made from field trial samples, using carbon-13 isotype discrimination (CID; Farquhar and Richards, 1984) as proxy for TE. CID from dry leaf biomass typically has a high heritability in both C3 (wheat, Rebetzke et al., 2002) and C4 cereals (maize, Gresset et al., 2014) and has been significantly negatively correlated to TE in C3 species such as wheat (Farquhar and Richards, 1984; Condon et al., 1990) and rice (Impa et al., 2005). However, CID is expensive to measure and its association with TE is more complex in C4 species (Farquhar, 1983; Henderson et al., 1996; Hammer et al., 1997).

Lastly, leaf-level measurements of TE are also commonly measured, but they are point measurements in terms of both space (part of a leaf) and time (seconds), and are typically more variable than plant-level measurements of TE. Combined with the time-consuming nature of these measurements, they are generally less conducive to large-scale phenotyping than plant-level measurements, as suggested by the results from this study concordantly with previous reports (e.g. Chenu et al., 2018; Vadez et al., 2024).

### 4.3 Implication for the industry

In addition to the new proposed methodology for screen TE proxy, this study brings to breeding a list of new lines from diverse backgrounds which substantially outperformed benchmarking elite cultivars for TE and TR_norm-highVPD_ (Fig. 6,Fig. **7**). Those include Australian-adapted lines derived from a cross between two high-TE Australian cultivars (Drysdale and Suntop; Condon et al., 2004; Rebetzke et al., 2009; Tausz-Posch et al., 2012; Fletcher et al., 2018; Fletcher, 2020; Collins et al., 2021) and agronomically-sound diversity core collection lines (Balfourier et al., 2007) which may bring new alleles to Australian breeding pools. Some of these promising lines depicted a TR_norm-highVPD_ which almost plateaued for VPD≥1.3-1.5 kPa (Fig. 9). As such, those lines are expected to bring substantial potential yield benefits through a shift of water use, with early water saving allowing more efficient water usee during post-flowering critical stages, especially in environments where crop heavily rely on stored soil moisture (Collins et al., 2021; Borrell et al., 2023). However, such lines could suffer yield loss in years when water is not limiting, as reduced TR_norm-highVPD_ may restrict CO_2_ uptake and biomass accumulation (Collins et al., 2021). Interestingly, limited transpiration at high evaporative demand (e.g. TR_norm-highVPD_) has been a cornerstone in the success of the recently developed Pioneer AQUAmax maize hybrids in the US (Cooper et al. 2014, Messina et al. 2015), thus reinforcing the potential benefits from the current study.

Results from this study provide quality data to parameterise crop models as done by Collins et al. (2021) to quantify the potential value of existing genotypic variation. While limited normalised transpiration rate under high evaporative demand (e.g. TR_norm-highVPD_) has been identified as a valuable trait for various crops and production regions (e.g. Sinclair et al., 2005, 2010; Messina et al., 2015), quantifying the value of this trait based on ‘real’ genetic variations as rarely been done (Collins et al., 2021). Further, in the current study, two types of promising genotypes for TE were identified: some outperformed modern benchmark cultivars for biomass production while using similar amount of water, while others accumulated similar biomass than elites while using less water (Fig. 7). Unravelling the value of these strategies for the current and projected target populations of environments (TPEs) would inform breeding. Crop models also open further avenues to predict genotype × environment × management (G×E×M) interactions that influence the expression and utility of those traits singly and/or in combination with other adaptive traits (Hammer et al., 2014; Durrington et al., 2025) in relevant management systems (e.g. Zheng et al., 2018) for current and future climates (Lobell et al., 2015; Chenu et al., 2017; Collins and Chenu, 2021).

## 5 CONCLUSION

This study demonstrates that reduced normalised transpiration under high evaporative demand (TR_norm-highVPD_) is a key driver of genotypic variation in TE in wheat. This trait strongly correlated with TE, depicted stable genotype ranking across environments, and was evident in diverse genetic backgrounds. Several high-TE low-TR_norm-highVPD_ lines, including Drysdale × Suntop derivatives and core collection accessions provide promising candidates for (pre-)breeding.

The framework developed here offers a scalable and physiologically grounded approach to screen for high TE, using plant-level transpiration responses to environmental demand without relying on destructive measurements or approximative estimation of leaf area or biomass. It also allows trait screening in fluctuating atmospheric conditions similar to production environments, and with similar plant densities mimicking production systems.

The study offers plant breeders a robust phenotyping target and methodology to enhance selection for drought adaptation and provide researchers with trait-based parameters that can be directly integrated into crop models and genomic selection pipelines.

While the potential values of TR_norm-highVPD_ and TE have been demonstrated experimentally and *in silico*, trade-offs of these traits can limit productivity in non-limiting water conditions, e.g. as reduced transpiration may constrain biomass accumulation when moisture is ample. As such, the integration of TR_norm-highVPD_ and TE into breeding programs should be supported by crop modelling and multi-environment testing to ensure yield advantages across diverse production systems.

## ABBREBIATIONS

LTR: limited transpiration rate at high evaporative demand
TE: Transpiration efficiency
VPD: vapour pressure deficit
DS lines: lines derived from a cross between Drysdale and Suntop
g_s_: stomatal conductance
NIL: near isogenic line
TR_norm_: normalised transpiration rate
TR_norm-highVPD_: normalised transpiration rate at VPD above 1.3 kPa
TR7_GLAnorm-highVPD_: cumulative transpiration over the last 7 days of the season normalised by green leaf area at harvest
WUE: water use efficiency

## 6 SUPPLEMENTARY DATA

Supplementary data are available at JXB online.

7 ACKNOWLEDGEMENTS

The authors thank John Evans for his advice on gas exchange measurements, Matthew Reynolds, Jack Christopher, and Penny Tricker for providing genetic material, Greg McLean, Bangyou Zheng, Claire Harris, Najeeb Ullah, Jingke Liu and Hannah Collins for their technical assistance, Scott Chapman and Bangyou Zheng for access to the controlled environment facility, and Graeme Hammer for valuable discussions. This research has been funded by the Australian Research Council (ARC) Centre of Excellence for Translational Photosynthesis, the ARC Linkage Project LP210200723 and the Grains Research and Development Corporation (Project CSP00179).

## 8 AUTHOR CONTRIBUTIONS

**Brian Collins:** Data curation, Formal analysis, Investigation, Methodology, Project administration, Software, Visualization, Writing – original draft.

**Karine Chenu:** Conceptualization, Formal analysis, Funding acquisition, Methodology, Project administration, Resources, Supervision, Validation, Writing – review & editing.

## SUPPLEMENTARY MATERIAL

**Table S1.**
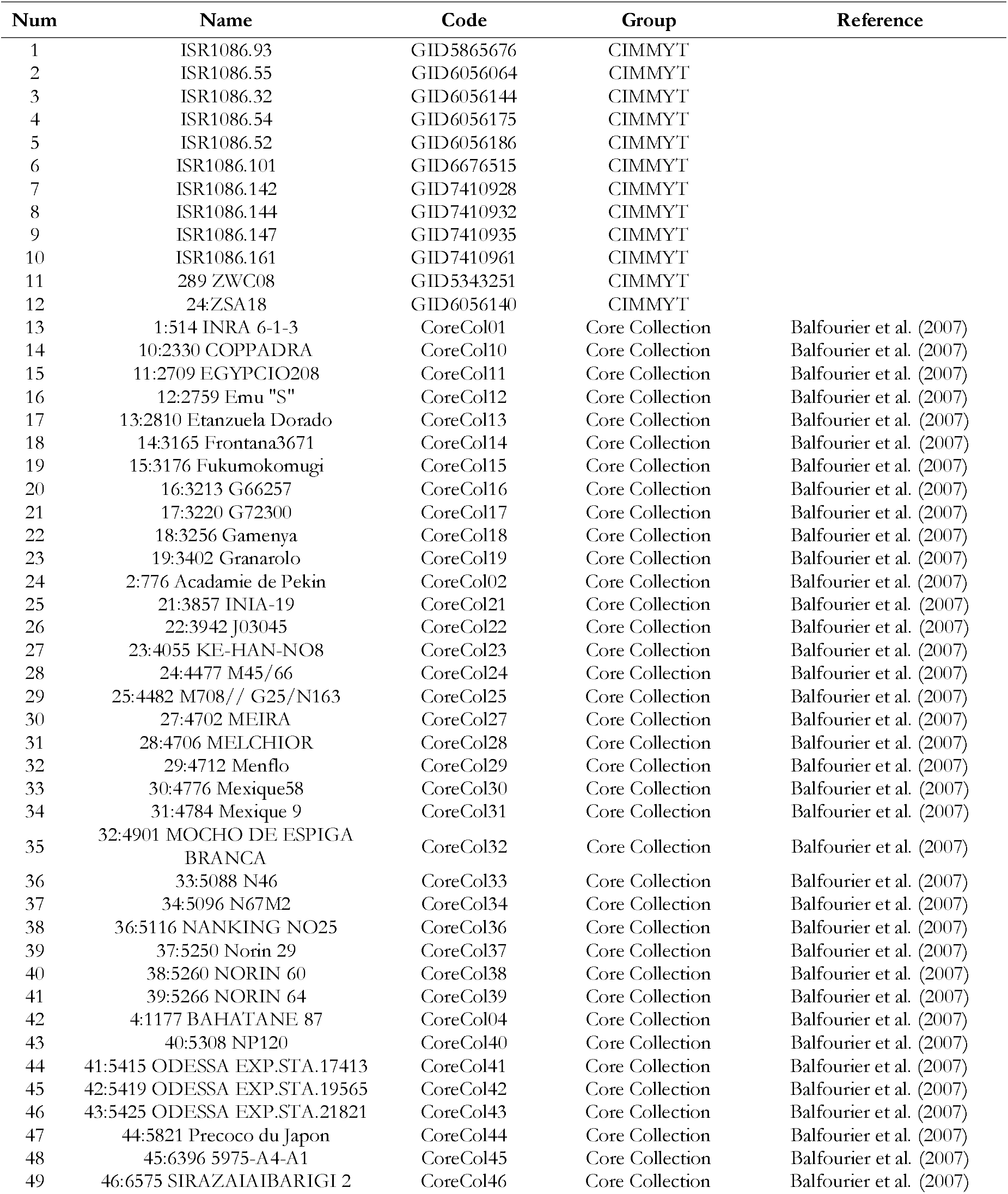
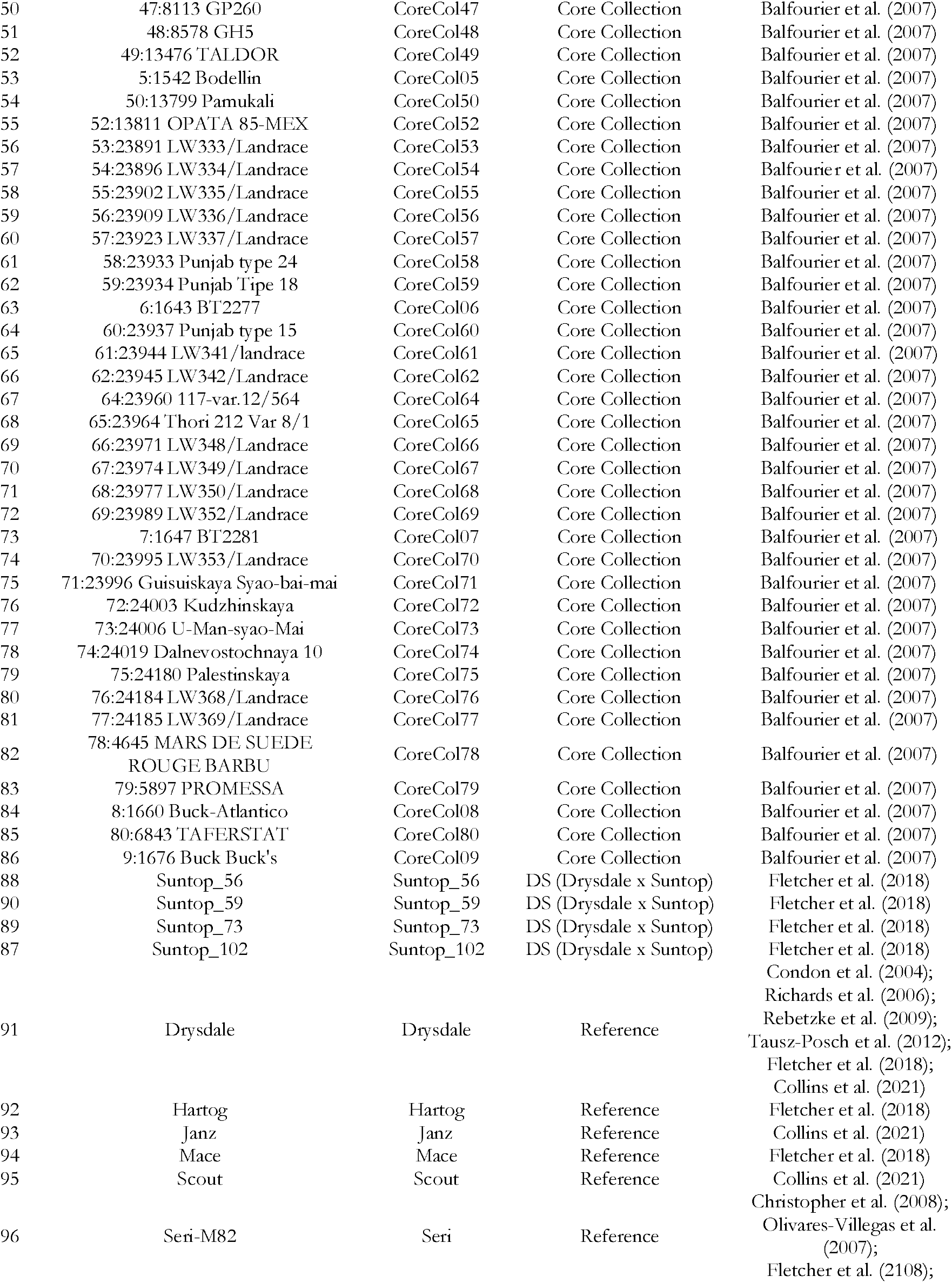
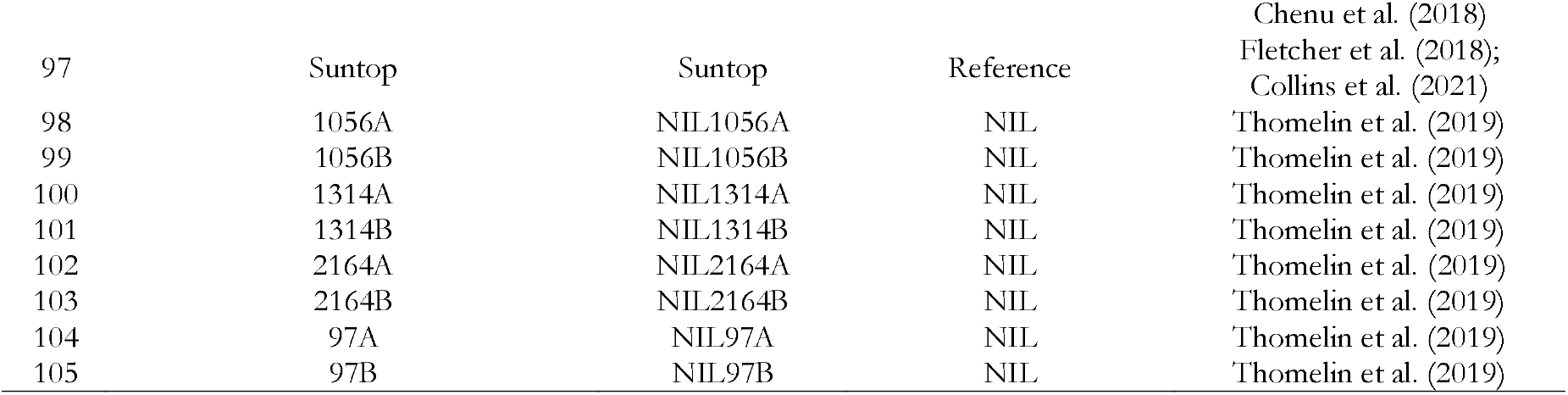
List of the 105 wheat studied genotypes.

**Fig. S2.**
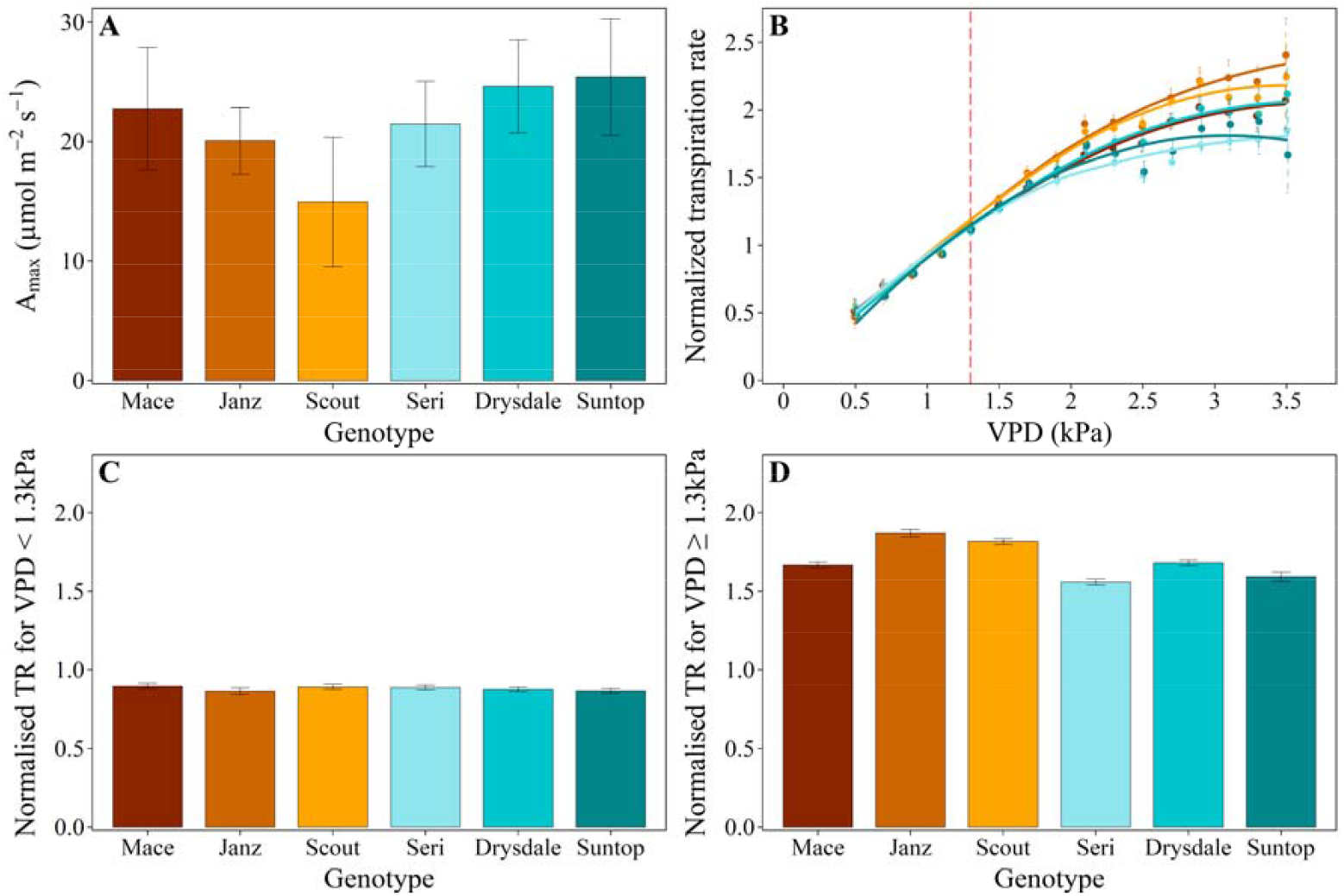
Physiological traits associated with transpiration efficiency for six reference genotypes in 2019 (Exp 2): (A) maximum photosynthetic capacity (A_max_), (B) normalised transpiration rate (TR_norm_) response to VPD, (C) average TRnorm for VPD<1.3 kPa (TR_norm-lowVPD_), and (D) average TR_norm_ for VPD≥1.3 kPa (TR_norm-highVPD_). Error bars correspond to 95% confidence intervals. Data for Exp 1 (2018) are presented in Fig. 2.

**Fig. S3.**
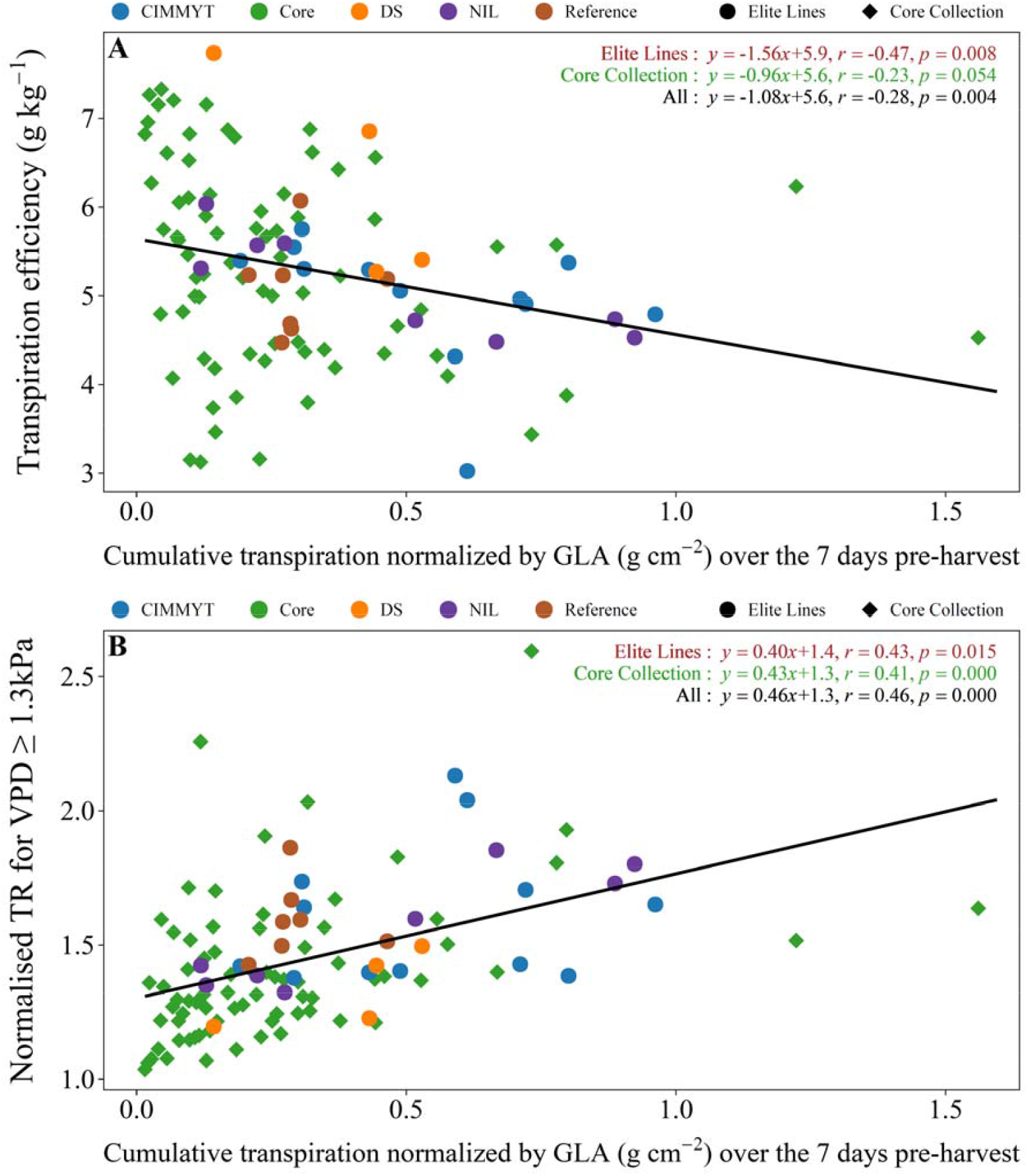
Relationship of (A) transpiration efficiency and (B) average normalised transpiration rate (TR) under high evaporative demand (TR_norm-highVPD_), against cumulative transpiration over the last 7 days (before harvest) normalised with green leaf area (GLA at harvest) for VPD≥1.3 kPa and Radn≥0.84 MJ m^-2^ h^-1^ (TR7_GLAnorm-highVPD_). Data from Exp 2. ‘Elite Lines’ include elite breeding lines from CIMMYT lines, NIL and DS lines (which have Australian cultivars as genetic backgrounds) and the reference cultivars. Refer to section 2.1 or the caption of Fig. 6 for the definition of the five groups of genotypes evaluated.

**Fig. S4.**
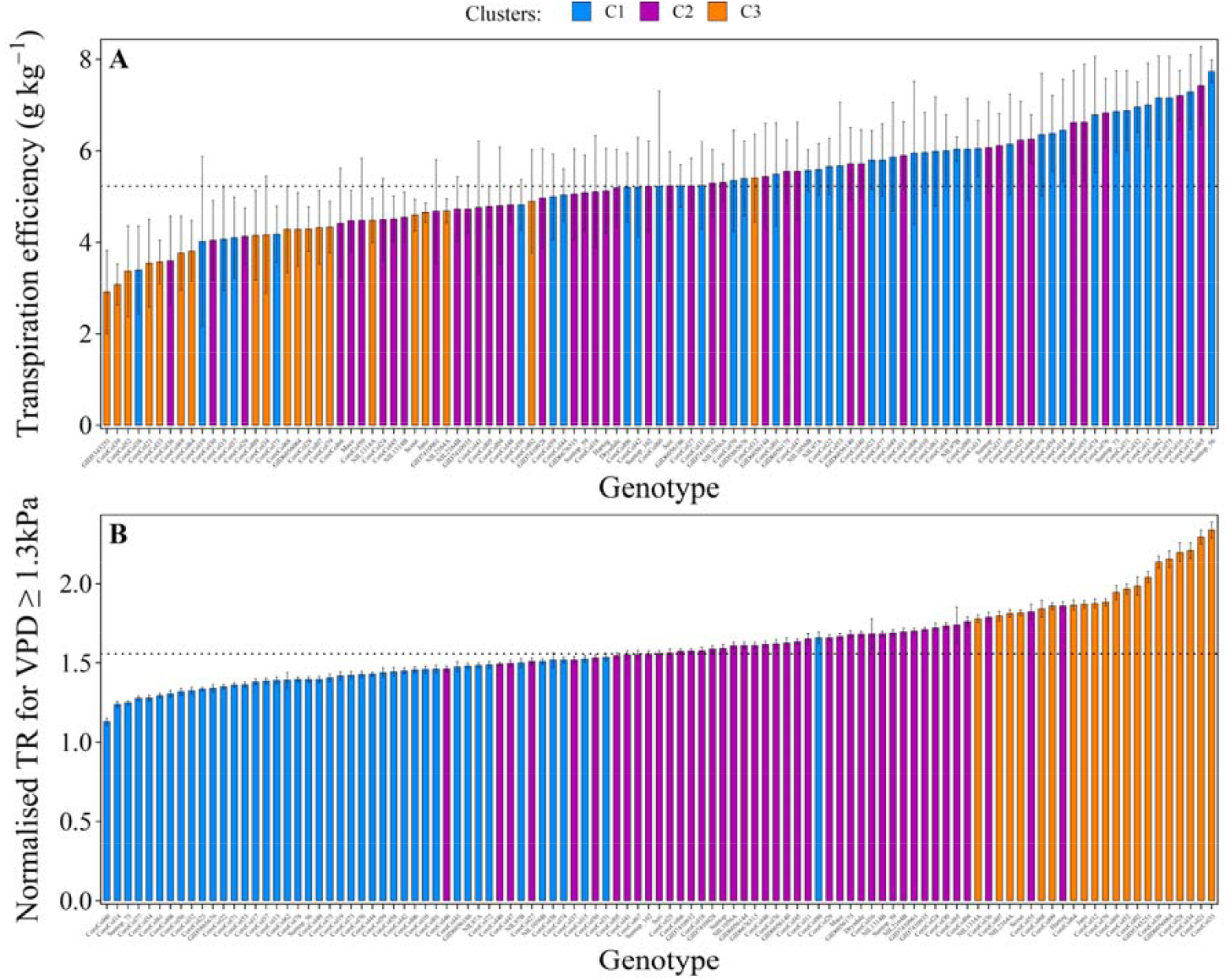
(A) Transpiration efficiency and (B) normalised transpiration rate at high evaporative demand (TR_norm-highVPD_) for all 105 genotypes coloured depending on their response to VPD (three clusters). Normalised transpiration rates correspond to genotypic averages for VPD≥1.3 kPa and Radn≥0.84 MJ m^-2^ *h*^-1^ between 750^°^Cd after sowing and harvest. Data from Exp 1 (2019). Clusters were defined based on their responses of normalised transpiration rate to VPD (Fig. 9B) and correspond to low (C1), medium (C2) and high (C3) TR_norm-highVPD_. Dotted lines show the median values in each panel. Error bars correspond to 95% confidence intervals. Refer to section 2.1 or the caption of Fig. 6 for the definition of the five groups of genotypes evaluated.

